# Basal forebrain activation improves working memory in senescent monkeys

**DOI:** 10.1101/2024.03.01.582925

**Authors:** Kendyl R Pennington, Luca Debs, Sophia Chung, Janki Bava, Clément M Garin, Fernando L Vale, Sarah K Bick, Dario J Englot, Alvin V Terry, Christos Constantinidis, David T Blake

## Abstract

Brain aging contributes to cognitive decline and risk of dementia. Degeneration of the basal forebrain cholinergic system parallels these changes in aging, Alzheimer’s dementia, Parkinson’s dementia, and Lewy body dementia, and thus is a common element linked to executive function across the lifespan and in disease states. Here, we tested the potential of one-hour daily intermittent basal forebrain stimulation to improve cognition in senescent monkeys, and its mechanisms of action. Stimulation in five animals improved working memory duration in 8-12 weeks across all animals, with peak improvements observed in the first four weeks. In an ensuing three month period without stimulation, improvements were retained. With additional stimulation, performance remained above baseline throughout the 15 months of the study. Studies with a cholinesterase inhibitor produced inconsistent improvements in behavior. One of five animals improved significantly. Manipulating the stimulation pattern demonstrated selectivity for both stimulation and recovery period duration. Brain stimulation led to acute increases in cerebrospinal levels of tissue plasminogen activator, which is an activating element for two brain neurotrophins, Nerve Growth Factor (NGF) and Brain-Derived Growth Factor (BDNF). Stimulation also led to improved glucose utilization in stimulated hemispheres relative to contralateral. Glucose utilization also consistently declines with aging and some dementias. Together, these findings suggest that intermittent stimulation of the nucleus basalis of Meynert improves executive function and reverses some aspects of brain aging.

**Highlights:**

- The basal forebrain and its cholinergic projections are the sole source of acetylcholine for the cortical mantle in primates and humans.
- Forebrain function tracks cognitive loss throughout the adult lifespan.
- One hour per day intermittent stimulation of this region improves executive function behaviors and plausibly reverses some aspects of brain aging, a large risk factor in dementias.
- This stimulation exceeds impacts of standard pharmacotherapies, is enduring, recruits brain neurotrophic pathways and improves cortical glucose utilization.

## Introduction

The cerebral cortex of mammals receives innervation from forebrain cholinergic neurons(1), which facilitate neurotransmission and maintain blood flow and neurotrophic tone in the neuropil(2, 3). The neurodegeneration in this pathway parallels declines in cognition that occur with aging and dementia, and in early phase degeneration in Alzheimer’s disease(4–8). The cholinergic basal forebrain region in the inferior aspect of the globus pallidus is the nucleus basalis of Meynert in primates, and it supplies cholinergic innervation to the cortical mantle(9).

The aged monkey represents a reasonable model to investigate the potential of deep brain stimulation of nucleus basalis of Meynert to halt or reverse cognitive decline. The Rhesus macaque ages at roughly three times the rate of humans(10). The aged primate has declines in cognition analogous to those in humans(11). Beta-amyloid plaques form with age although the presence of neurofibrillary tangles composed of tau is more rare and appears shifted to a later age(12). We investigated whether deep brain stimulation of the nucleus basalis of Meynert in senescent monkeys could halt or reverse cognitive decline.

## Results

Sixteen Rhesus monkeys (*Macaca mulatta*) over the age of 25 (analogous to age >75 in humans) underwent stimulation and/or pharmacological administration in these experiments. We evaluated the behavioral effects of stimulation primarily in five Rhesus monkeys (2 male, 3 female) that were implanted bilaterally with stimulating electrodes and had stable behavioral performance. These monkeys completed sufficient numbers of daily trials before and after stimulation to quantify stable delay period thresholds in an adaptive tracking paradigm (Figure 1B) version of a working memory task (Figure 1A). Prior to implantation, animals learned the task requiring them to remember a cue stimulus presented at the center of a screen. After a variable delay period, the monkey selected between two test stimuli, one of which matched the cue. The behavior featured a two-alternative forced choice procedure and used adaptive tracking to adjust the duration of the delay period. If the monkey chose correctly on three consecutive trials, the trial delay grew roughly 50% longer. Each error resulted in the delay being shortened by 33%. This 3-up, 1-down adaptive tracking can be analyzed by averaging an even number of tracking reversals to converge on the 79% correct delay threshold(13) and allowed monkeys to perform using the same task parameters daily. Our primary behavioral metric was the mean of adjacent pairs of adaptive tracking reversals, one upward facing, and one downward facing, which has the expected value of the 79% correct threshold.

**Figure 1.**
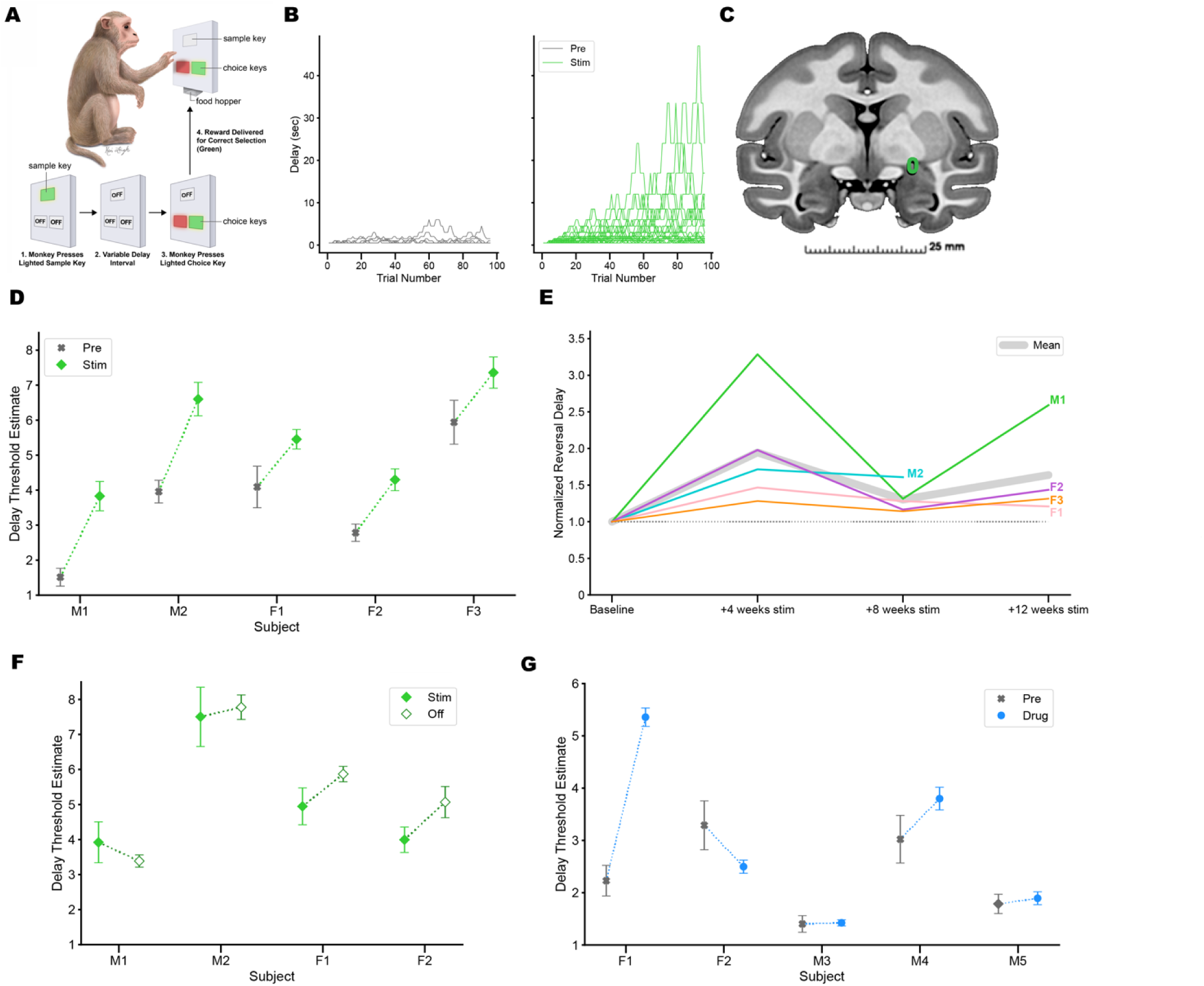
A. Monkey task. The monkey touches the cue on the touchscreen. The squares darken for the delay period. Then, a match and a distractor appear in the lower squares for the choice. B. Adaptive tracks for monkey performance for one animal before stimulation onset (left), and in the last four days of stimulation in the first 12 weeks (right). Each line traces the delays used in one training session. C. Target electrode placement from CT superimposed on a coronal MRI. The target was the mediolateral center of the globus pallidus, in the floor of the globus pallidus, 2 mm posterior to the crossing of the pallidus by the anterior commissure. MRI section was taken from the NMT template V2 (20). Green outline depicts the activating function radius of the implanted electrode. D. Single animal adaptive tracking threshold estimates in the control period and the ensuing first 8-12 weeks of stimulation. E. Normalized individual changes in threshold each four weeks, and group average (thick gray line), for the same five animals as in D. F. Threshold changes compared across the last four weeks of stimulation for four of the animals in D compared to the next 12 weeks after ceasing stimulation. G. Changes caused by donepezil treatment for 12 weeks in five animals not receiving stimulation.

The implantation target was the floor of the globus pallidus, in the medial-lateral center of the pallidal floor, and 2 mm posterior to the crossing of the globus pallidus by the anterior commissure. Placement was verified with CT imaging, shown in Figure 1C and Figure S1, and/or histological lesions, postmortem. After implantation and recovery, animals performed the working memory task daily, and received unilateral deep brain stimulation consisting of a 60 Hz train of pulses for 20 seconds interspersed with 40 second periods without pulses, for one hour daily. A pulse amplitude of 0.5 mA was chosen to produce an estimated activating function radius of 700 μm. We were particularly interested in the usefulness of deep brain stimulation as a means of improving cognitive function even when not applied concurrently with task execution. Therefore, we delivered stimulation during the task on half the sessions, the during condition, and delivered the stimulation after the task on half the sessions, the after condition. All data in Figures 1 and 2 were recorded in the after condition.

**Figure 2.**
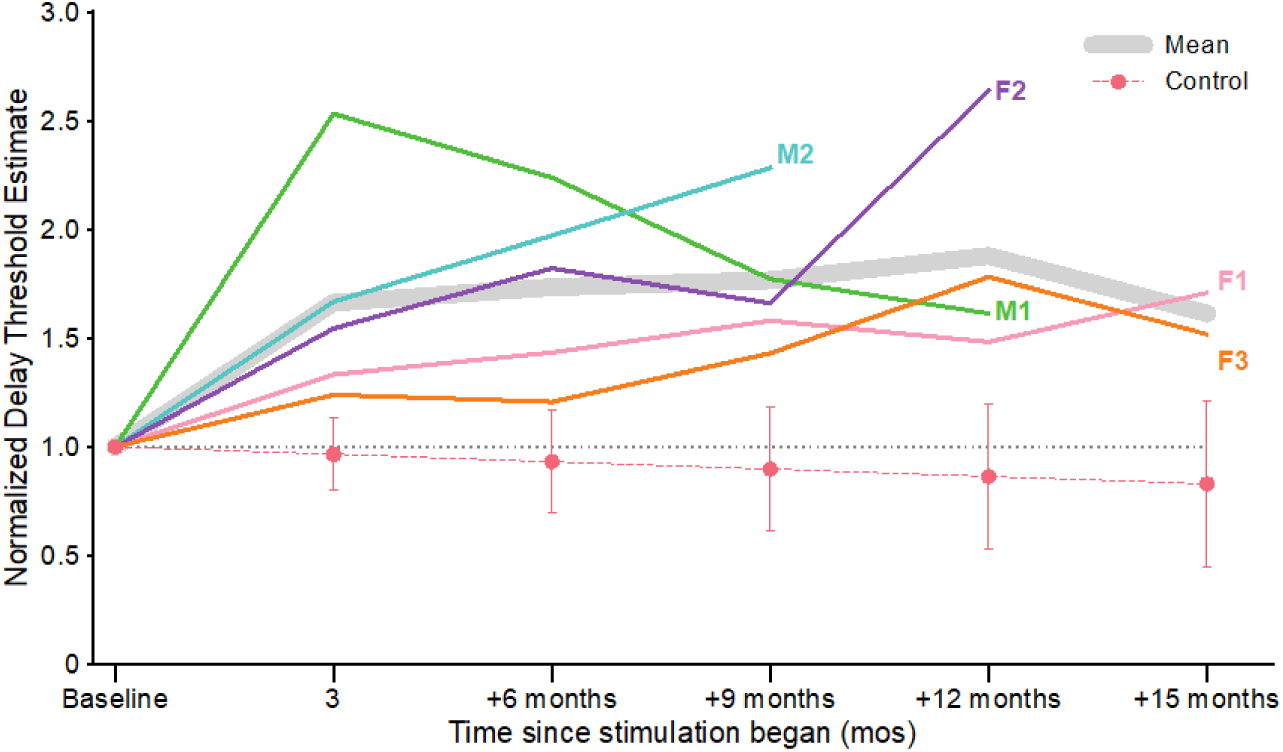
Long-term changes in the five animals from Figure 1D. The three month time period includes the three month average data from Figure 1D, while the other time points include data from 1F and after in which variable stimulation protocols were followed as detailed in Figure S6. The control data are shown lower in the figure with error bars measuring one standard deviation. The dotted line shows unity.

### Cholinergic DBS produces long-term cognitive improvement

Over the three-month period after the initiation of stimulation, the animals’ delay thresholds were higher (Figure 1B, D). In this 12 week period, the monkeys were correct more often, and tracked to lonqger delays. Their changes in threshold are shown in Figure 1D. A generalized linear mixed-effects model (GLMM) in which monkey identity was treated as a random effect and stimulation was a fixed effect estimated a 49% increase in threshold, which was significant (main effect of treatment, Z = 6.83, p=7.3e-12). Post hoc t-tests indicated that each of the five monkeys significantly increased their delay threshold (M1 t_33_=4.32, p=0.00013, M2 t_99_=3.34 p=0.0012, F1 t_33_=2.42, p=0.022, F2 t_135_=3.48, p=0.00068, F3 t_98_=3.05 p=0.0029).

To assess the time course of changes, we grouped the monkey threshold estimates in four week periods after the initiation of stimulation. These are plotted after normalization in Figure 1E. A GLMM using each four week period as a fixed effect found that the 0-4 week period was 34% larger than the 4-8 week period (main effect of treatment, Z=3.80, p=7.2e-5), and the 8-12 week period was 19% larger than the 4-8 week period (main effect of treatment, Z=2.30, p=0.026). Individual animal thresholds were all highest in the first four weeks, which indicates we observed the largest changes from stimulation in those first four weeks.

In the three months after the initial stimulation period, four of the monkeys were continued in testing for 12 weeks without further stimulation. All four subjects’ performance remained elevated above baseline in the off period, shown in Figure 1F, which suggests endurance of the long-term benefits of stimulation. A GLMM assessing stimulation stopping as a fixed effect resulted in a 14.4% increase in delay threshold after stimulation ended (main effect of treatment Z=2.385, p=0.017). Three monkeys had mean thresholds slightly higher in the 12 week off period than in the last four weeks of stimulation, and one had a slightly lower mean threshold, and no individual animals reached statistical significance in post hoc testing.

To further quantify effects over longer time periods, the animals that had received deep brain stimulation were given more stimulation, either unilateral or in some cases bilateral (see Figure S6), and their working memory thresholds tracked, for months after onset. All five animals retained above baseline working memory thresholds throughout study, over time periods as long as 15 months after implantation, as shown in Figure 2. A GLMM using stimulation as a fixed effect and comparing the pre-stimulation period to each three month period was significant for each three month period (Z scores 5.271, 6.162, 6.031, 6.669 at 6, 9, 12, and 15 months). The three month data points in Figure 2 are the same depicted in Figure 1D.

Control monkey data, which consisted of animals in the same age range that performed the same task over months of time without stimulation, were compiled for comparison. Working memory performance declines with age in macaques, with animals over age 25 reaching delays that are approximately 50% lower than animals under age 10 (14). Each animal’s threshold was measured over each nonoverlapping 90 day period for which data existed, and then averaged within each animal to create a distribution from these nine animals. The average three month change across these nine control animals was a 3.4% decrease in working memory threshold, with a standard deviation of 16.6%. We projected this three month statistical trend each three months to fifteen months for comparison. At three months, the smallest difference in working memory delay change between the control mean and one of the five stimulated animals was a Z-score of 1.645. Five out of five animals exceeding that Z score has an associated probability of 3.1×10^-7^. At the 6, 9, 12, and 15 month intervals, the data were also significant (minimal Z=1.17, n=5, p=2.59×10^-5^, Z=1.857, n=5, p=3.18×10^-8^, Z=1.865, n=4, p=1.12×10^-6^, Z=1.65, n=2, p=0.002). These sample control data are shown below the stimulation animal data in Figure 2.

### Cholinesterase inhibitors and off-target stimulation

To assess whether cholinesterase inhibitors could also provide enduring improvement of working memory, monkeys were given the commonly prescribed cholinesterase inhibitor donepezil daily after their working memory tests. The treatment of dementia often applies donepezil, which prolongs the action of acetylcholine by inhibiting its hydrolysis. We have previously shown that it can acutely improve working memory in monkeys if given before testing(15). Monkey performance in the period before and after donepezil administration is shown in Figure 1G, for a cohort of five monkeys that did not receive stimulation during this testing period. A GLMM on the impact of donepezil on delay threshold resulted in a significant 15.9% increase in delay threshold (Z=2.378, p=0.017). Post hoc t-tests found a positive significant impact only in animal F1 (t_28_=7.29, p = 6.02e-8). Animals that appear both in Figure 1D and 1G were tested for the impact of donepezil prior to washout and implantation.

A further positive stimulation control treatment consisted of electrodes that were not placed in the subpallidal basal forebrain and were stimulated with the same parameters as the treatment group. As shown in Figure S3, this treatment did not result in a consistent improvement in working memory duration. Electrode misplacement by more than 2 mm from the desired targets occurred in either the mediolateral or anterior posterior plane. A GLMM using stimulation as a fixed effect on these three animals resulted in a significant 17.0% decrease in threshold caused by stimulation (main effect of treatment, Z=2.46, p=0.014).

### Timing and duration of stimulation

Prior work has shown that basal forebrain stimulation could acutely impair or improve working memory depending on the stimulation pattern in young monkeys(15, 16), in addition to producing long-term improvements. We examined this effect in elderly monkeys by comparing behavior in the during condition to the after condition, shown in Figure 3A. A GLMM using acute stimulation as a fixed effect, as opposed to interleaved behavioral days in the after condition, revealed that thresholds were 12.9% lower during stimulation (main effect of treatment, Z=2.784, p=0.005). Post hoc testing reached significance for lower thresholds in the during condition only for monkey M1 (t_253_=4.38 p=0.00001). Four of the five animals had lower mean and median threshold estimates in the during condition, while one had a lower mean and median condition in the after condition. This effect contrasts with our observations in young monkeys using the same stimulation pattern, 20 seconds on and 40 seconds off, for which all animals performed better in the during condition. We next examined whether these elderly monkeys would benefit from different stimulation patterns.

**Figure 3.**
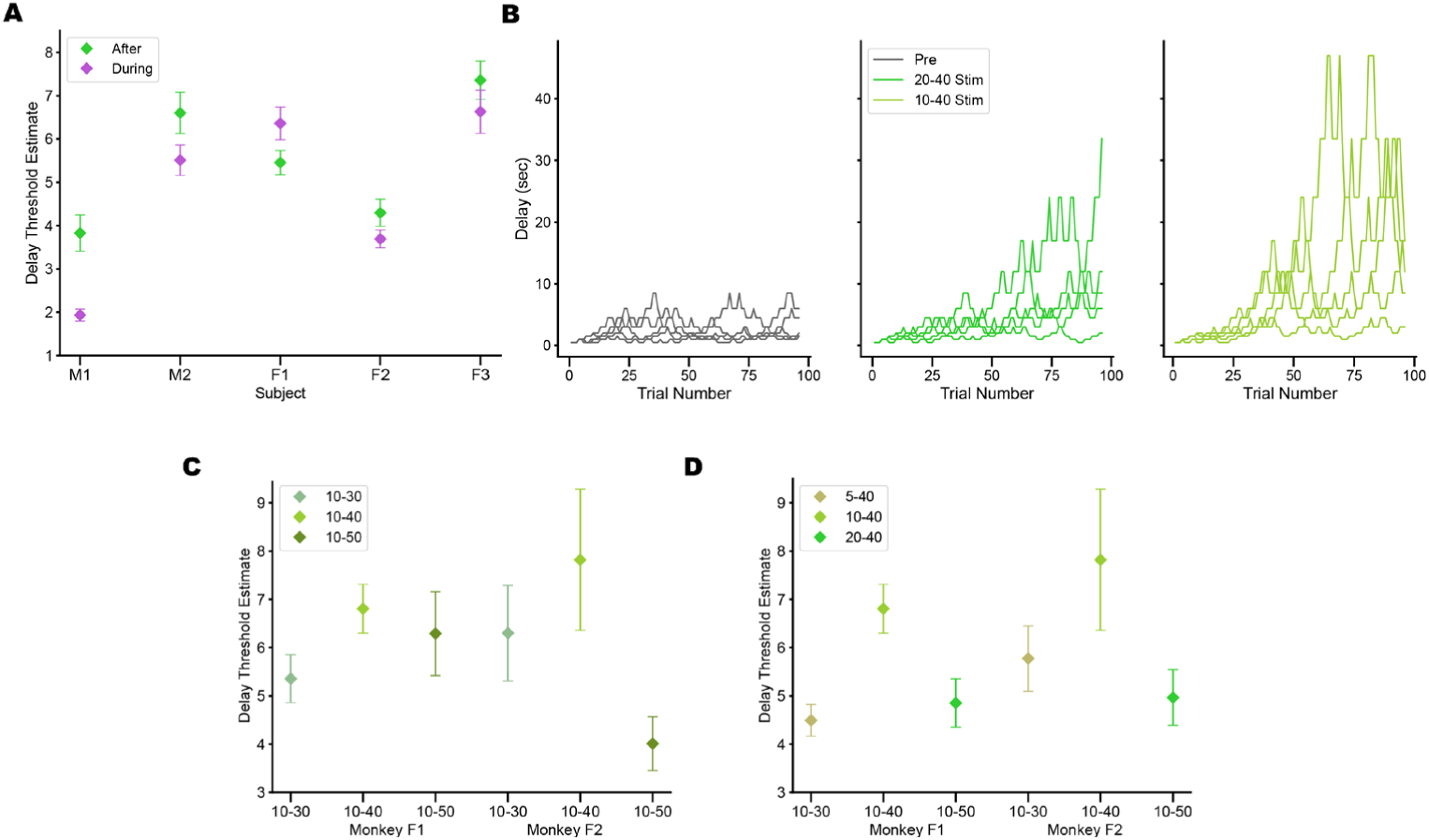
A. Acute effects of stimulation. Plotted in purple are estimates of threshold taken during stimulation, and plotted ingreen are estimates of threshold on stimulation days in the after condition. Green data points represent the same data shown Figure 1D. B. Adaptive tracking delay plots from five sample behavioral sessions from monkey F2 during the prestudy period (left), during bilateral 20-40 stimulation (middle), and during 10-40 stimulation (right). C. Threshold estimate distributions for animals F1 and F2 under different bilateral recovery period conditions. D. Threshold estimate distributions for animals F1 and F2 under different bilateral stimulation period conditions.

The stimulation pattern was altered to use a balance of stimulation and recovery that had less stimulation or more recovery. The prior parameters had impacts on long-term behavior that were found serendipitously, and not through systematic exploration of the parameters. Recent mouse imaging work also suggested that the stimulation period was too long, and the recovery period too short, to achieve maximal acetylcholine bursts in each cycle(17). The two animals most recently implanted in our study (subjects F1 and F2) were stimulated bilaterally, and different parameters were tested.

As shown in Figures 3C-D, S4, and S5, working memory threshold was highest during periods in which the stimulation parameters included a 10 second on period and a 40 second off period (10–40) in both animals. Example adaptive tracking delays from female monkey F2 are shown in Figure 3B for the pre-stimulation period on the left, after applying the 20-40 stimulation pattern in the center, and after applying the 10-40 stimulation pattern on the right. Two different experiments compared the effect of altering the stimulation period by comparing 5-40, 10-40, and 20-40 patterns, and of altering the recovery period by comparing 10-30, 10-40, and 10-50 patterns. In both cases, stimulation occurred at 60 pulses per second, and 60 cycles of the pattern were repeated. A GLMM assessing a fixed effect of stimulation period found a significant 28.4% longer delay threshold in the 10-40 period than in the 20-40 period, which was significant (main effect of treatment, Z=2.815, p=0.005). The difference between 10-40 and 5-40 was not significant, and the 10-40 mean was higher. A GLMM using the recovery period as a fixed effect found a significant 42.8% longer delay threshold in the 10-40 period than in the 10-50 period, which was significant (main effect of treatment, Z=2.815, p=0.006). The difference between the 10-30 and the 10-40 period was not significant, but the 10-40 mean was higher. The entire series of tested stimulation parameters are shown in Figure S4.

### Metabolic and neurotrophic mechanisms of long-term improvement

We investigated the underlying mechanisms through which stimulation improved cognitive performance in two ways. In the first, brain glucose utilization was evaluated in three animals by applying Fluorodeoxyglucose F18 (FDG) PET imaging before and after a 7–11-month period that included 2-3 months of unilateral stimulation. Glucose utilization is a marker that consistently tracks progression of aging and Alzheimer’s dementia(18, 19). We anticipated that the effect of time would be to decrease glucose utilization, while perhaps unilateral stimulation would reduce that decrease, or even increase utilization. Animals received one hour of daily intermittent unilateral stimulation, and the effect of stimulation on the standardized uptake value ratio (SUVr) relative to the ipsilateral pons was calculated(20). Both sides decreased in their average SUVr. The control side, contralateral to stimulation, averaged a decrease of 5.84% with a standard deviation of 0.70%. The stimulation side decreased by 3.59% with standard deviation of 2.12%. To look at potential regional changes in glucose uptake, a voxel-wise cluster analysis was performed. Starting with the control side (Figure 4A-left), the statistical maps were thresholded to identify contiguous clusters in which voxels trended down more strongly. These clusters with decreased utilization made up 18% of the hemisphere, and there were no clusters of increased utilization. Then, the stimulated side was analyzed after adjusting for the average changes in the control hemisphere. Three significant patches of relatively increased glucose utilization resulted (all t_8_>3.355, p<0.005, see Methods), with no clusters of decreased utilization, shown in Figure 4A-right. Using atlas parcellation from the precise CHARM atlas (AFNI)(21), we also compared glucose utilization cross-hemisphere by region in these three animals, and the regions strongly trended to have a smaller decrease in utilization on the stimulated hemisphere (40 of 51 comparisons positive, p = 5.70×10^-5^, binomial test) as shown in Figure 4B. Similar results were obtained when the cerebellum was used for the calculation of the SUVr (Figure S5, 38 of 51 comparisons positive, p=0.0003 binomial test).

**Figure 4.**
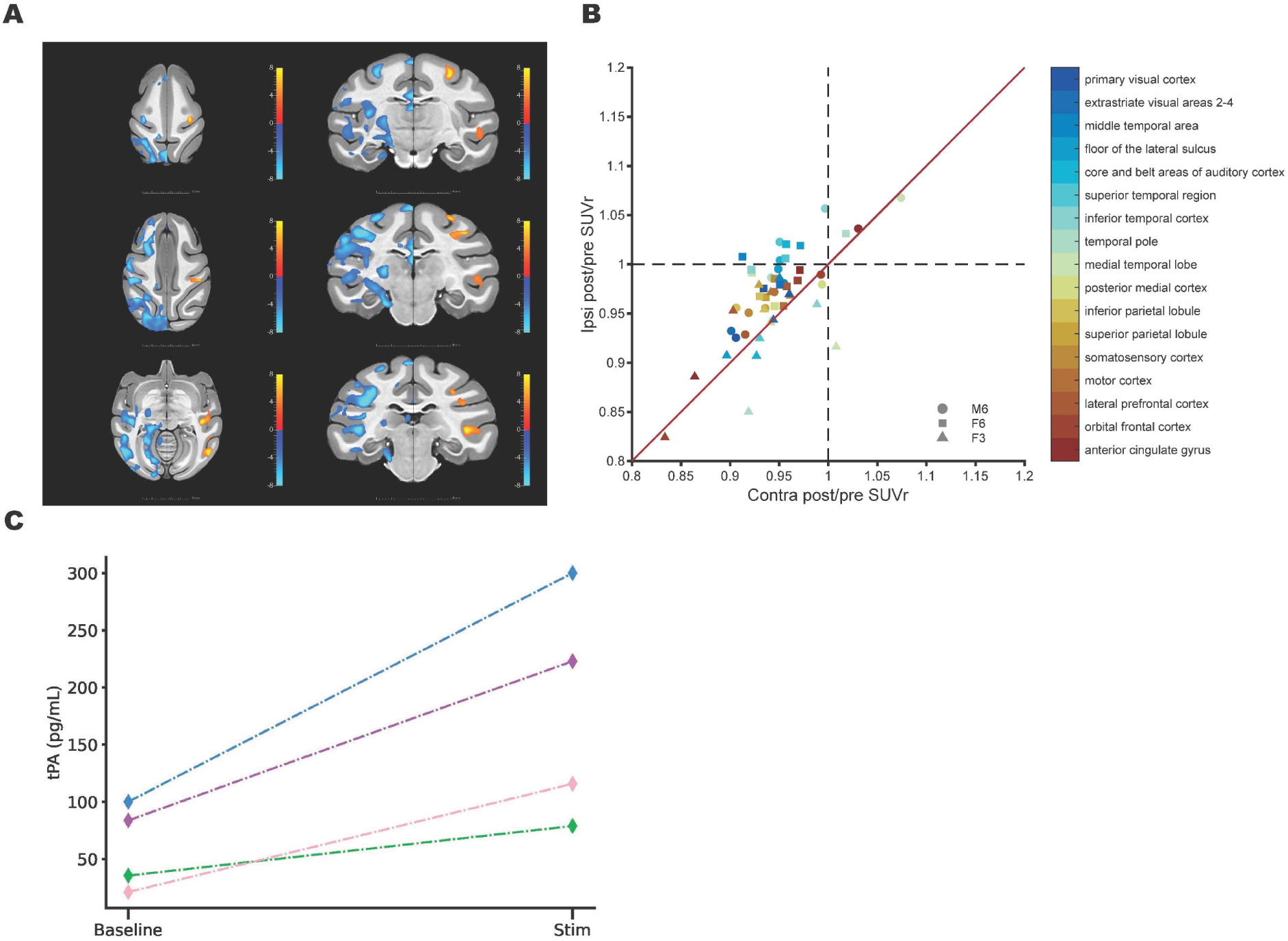
4. A. Statistical maps of lateralized effects on 18-F PET-FDG SUVr. Hemisphere on image left: significant clusters with time-dependent changes in glucose uptake. Hemisphere on image right: significant clusters with stimulation-dependent changes in glucose update, after controlling for effects in the control hemisphere. B. Comparison of change ratios in PET SUVr between stimulated and unstimulated in 17 cortical regions. C. Tissue plasminogen activator was measured in cerebrospinal fluid under neutral conditions or after a one hour stimulation period in four animals.

The second investigation of potential mechanisms underlying effects on working memory assessed whether activation of neurotrophic pathways was occurring in our preparation. Reduced neurotrophin expression has been documented in parallel with brain changes in aging and dementias, and increases in neurotrophins are being tested as treatments for dementias(22). Rodent work has indicated that expression of the receptors for the neurotrophins nerve growth factor (NGF) and brain derived neurotrophic factor (BDNF) are increased by this stimulation(17), which suggests that neurotrophins are being released and/or activated. This release is hypothesized to occur into the interstitial fluid of the cerebral cortex, which slowly mixes with cerebrospinal fluid(23). The fluid turns over in 6-8 hours, and mixes completely with cerebrospinal fluid (CSF) in less than half that time. In four of the monkeys, CSF was sampled before and after one hour of stimulation in the 20-40 condition. The fluid was tested for its concentration of tissue plasminogen activator (tPA), which is an activating element in the pathways for both NGF and BDNF. We found tPA levels in the same range as is documented in humans(24), and that the levels increased significantly and by up to three-fold by stimulation (t_3_=3.59,p<0.04), as shown in Figure 4C.

## Discussion

Deep brain stimulation has been used extensively in the treatment of movement disorders, and increasingly for the treatment of depression, epilepsy, and obsessive-compulsive disorder(25–27). Multiple studies have attempted stimulation of the nucleus basalis of Meynert in clinical trials(28–31). In most applications, continuous, 24 hour per day high frequency pulse trains are used to normalize the hyperactive circuits that receive the stimulation. Multiple hypotheses exist on the mechanisms that support this normalization(32). In contrast, shorter pulse trains are used in neurobiology to activate cortical circuits(33–35), rather than to suppress them. Limited duration pulse train stimulation of the basal forebrain increases cortical acetylcholine levels(36). Pulse stimulation depolarizes membrane potentials in proportion to the second spatial gradient of the voltage field, often termed the activating function(37). With suprathreshold, repetitive pulse stimulation, axons within the activating function radius will depolarize and generate action potentials(38), recruit closely synaptically coupled neurons(39, 40), and activate the local circuit.

We thus applied short pulse trains, with recovery periods, over one hour daily with intent on increasing the activity of the circuits. Our results establish this as a viable strategy for the long-term cognitive function improvement in aged individuals. Basal forebrain stimulation was superior to donepezil administration, and further, the effect was selective for the targeted subpallidal basal forebrain, as nearby, but off target, stimulation was impairing. The superiority of basal forebrain stimulation was evident in the overall increase in delay threshold, 49% for stimulation versus 15.9% for donepezil, and by the number of animals that showed a statistically significant improvement upon post hoc analysis, all five animals for stimulation and only one out of five animals for donepezil. This response rate for donepezil in aged monkeys is somewhat lower than the rate in Alzheimer’s clinical trials, although a wide range of response rates have been observed(41). The neurodegeneration of the basal forebrain region predicts non-motor executive function deficits in Alzheimer’s, Lewy Body and Parkinson’s dementia(7, 8, 42, 43), so its improved function from intermittent deep brain stimulation is a candidate for improving cognition in these populations.

The main mechanism of long-term improvement did not require stimulation to be delivered concurrently with behavior. The duty cycle of 20 seconds on, 40 seconds off led to impaired acute performance in aged monkeys. Our prior work in young male monkeys found consistent acute improvement with the same parameters(15, 16, 44). Together, these data suggest that the acute impact of intermittent stimulation using parameters defined from young male monkeys result in different neuromodulation in elderly animals. The tuning of the parameters for optimizing long-term improvements in working memory further revealed that half as much stimulation per cycle, 10 seconds instead of 20, resulted in 28.4% longer working memory duration. Elderly animals may require a different balance of intermittent stimulation and recovery to optimize acute and long-term effects.

Longitudinal PET-FDG imaging was used to examine the impacts of stimulation on brain glucose utilization. Generally, advancement in age is associated with decreased glucose utilization(45). The effect can be translated as a 7% loss in glucose utilization per decade of life(46, 47), and these findings apply equally in longitudinal scans of the same individuals. Our findings indicate that glucose utilization was decreased overall as expected in each cortical hemisphere compared to control regions not normally impacted by age or AD (i.e., the pons and cerebellum), but the decrease was lower in the stimulated hemisphere relative to the unstimulated hemisphere.

As intended, the stimulation perfuses the cerebral cortex with acetylcholine, with a recovery period, prior to perfusing the cerebral cortex again. The increased NGF and BDNF receptor expression found from this stimulation in mice(17), along with the increased CSF concentration of tPA observed in the present study, support the premise that this pattern of stimulation increases the forebrain projection-related acetylcholine release to enhance neurotrophic activity. Cholinergic activation has previously been shown(3, 48) to cause the release of neurotrophins and tissue plasminogen activator (tPA), which lead our hypotheses for underlying processes causing the improved behavioral function. The neurotrophins are necessary for multiple forms of synaptic and map plasticity in the cerebral cortex(49, 50), and in the nonhuman primate preparation tPA and neurotrophins may be readily measured by their concentrations in CSF. The release of NGF by stimulation should elaborate the arborization of cholinergic axons into the cerebral cortex, and result in greater acetylcholine release(51). BDNF, in turn, has effects that include increased coupling between parvalbumin interneurons and pyramidal neurons(52).

The potential application of this form of stimulation to improve executive function, especially in dementia, is also beginning to be explored. Three Parkinson’s patients, receiving this pattern of stimulation for two weeks, improved in their sustained attention, compared either to baseline, or to performance after two weeks of continuous pulsetrain stimulation(53). Deep brain stimulation to reduce activity in the stimulated neural circuits has been heavily explored for the past two decades; this work, and the work of others(35), are beginning to explore the potential of brain stimulation to increase activity in the stimulated neural circuits.

## Acknowledgments

We would like to thank K. Shanazz and R. Jagirdar for their histological help in localizing electrolytic lesions, C. Suell, K. Clemencich, K. Yetman, M. Plagenhoef and J. Sun for technical help with animal testing and surgeries and C. Cryan for work in the first ELISA tests of CSF for tPA. Keri Leigh Alber created the artwork in Figure 1A. Srikantan Nagarajan provided helpful feedback on the statistical approach for the behavioral analysis.

## Author Contributions

DTB and CC designed and planned this study. KRP and DTB adapted the behavioral apparatus of AVT that was used for all behavior. KRP performed all behavioral statistical analysis with supervision from DTB and CC. KRP designed and implemented the tPA ELISA work. Animal behavior was conducted by SC and JB (Vanderbilt) and KRP (Augusta). Surgical methods were created and implemented by LD, FV, DJE, SKB, DTB, and CC. LD and FV implemented ventricular drains for the tPA work. PET experiment design and analysis were performed by CMG and CC. Donepezil work was designed by AVT and implemented by KRP, SC, and JB. Manuscript was drafted by CC, DTB and KRP and edited by all authors.

## Data and Material Availability

all raw data will be uploaded and accessible freely upon publication. Implantable pulse generators and accompanying hardware were provided by Boston Scientific through a public/private partnership. Contact about any other materials.

## Supplementary Materials for

### Materials and Methods

Animals used in this study were 16 Rhesus monkeys Macaca Mulatta over 25 years in age. All had been used previously in behavioral pharmacology studies. Studies were performed at Augusta University (M1, M2, F1, F2, M3, M4, F4, F5 and others) and Vanderbilt University (F3, F6, M5, M6 and others). The code M in the monkey identification indicates a male, and a F indicates a female. All studies were approved by the IACUC at either Vanderbilt or Augusta University.

### Surgical implantation

Surgeries were performed under isoflurane anesthesia. A methods paper describes these procedures in detail(54). Animals were placed in a stereotaxic instrument using the ear canals and lower orbital ridges as fiduciary points. The Vanderbilt University animals had pre-operative MRI scans, and T1 images were used for planning targets. Augusta University used stereotaxic coordinates of 9 mm lateral, 19 mm anterior, and 29 mm deep from the lower surface of the bone. Similar coordinates were used at Vanderbilt except that the locations were corrected for altered alignment between the fiduciary points and the brain.

For the animals with imaging alignment, CT and MRI scans first were acquired. CT scans were acquired with a Philips (Amsterdam, Netherlands) Vereos PET/CT scanner, while MRI scans were acquired with a Philips 3T Ingenia Elition X scanner. MRI scans included a T1-weighted image with 0.5-mm isotropic voxels. The T1w image was skull-stripped and the brain parcellated into brain regions according to the symmetric NIMH Macaque Template (NMT) v2.0 of 0.25-mm isotropic voxels and the accompanying Cortical Hierarchy Atlas of the Rhesus Macaque (CHARM)(21) and Subcortical Atlas of the Rhesus Macaque (SARM)(55) using the @animal_warper function as part of the AFNI software(56). This also generated a set of linear and nonlinear transforms that map the subject spaces from and to the template space. The CT and T1w images were aligned using the General Registration (BRAINS) module in 3D Slicer(57) which finds and applies a rigid-body transform. Alignment quality was manually verified. Basal forebrain targets were identified from the atlas and visually confirmed/adjusted using the T1w image.

A Brainsight Vet system (Rogue Research Inc., Montréal, Canada) was used to plan the trajectory of implantation pre-surgically. The landscape of the skull surface was reconstructed from either the CT scan or the T1w image. The co-registration between the skull and the basal forebrain target allowed the determination of the electrode’s entry point and inserting angle.

The target was 2 mm posterior to the crossing of the globus pallidus by the anterior commissure, and in the floor of the globus pallidus. Bone screw holes 2-3 mm in diameter were drilled over the targets, and a bone screw with a hollow center (Gray-Matter Research, Bozeman MT) was installed. A guide cannula was inserted to point at the target. Electrodes were lowered into the cannula, the cannula was removed, and the electrodes were adjusted in depth. Silicone (Kwik-Sil, WPI Inc, Sarasota, FL, USA) was poured over the rear of the electrode to stabilize it in depth. After both electrodes were inserted, a wire connector from Boston Scientific, “a pigtail”, was tunneled between the scalp and subscapular space. Individual pigtail wires were resistance welded to the uninsulated rear end of the electrode wire. The connectivity of each wire to the pigtail connector was then checked. The welds were potted in silicone, and the pigtail was silk-sutured to the galea to prevent slip. The skin was closed with inverted simple interrupted absorbable sutures. A pocket was opened in the subscapular space, and the implantable pulse generator (Spectra Wavewriter IPG, Boston Scientific) was inserted. The IPG was not connected while welding was conducted. Thereafter, the IPG and pigtail were connected, and impedance was tested. Generally impedances under 8 kOhms were noted. The subscapular skin wound was closed as above.

### Electrodes

Electrode manufacture is described as in our methods paper(54). The electrode is a 0.003” diameter wire insulated with Teflon PFA (A&M Systems, Sequim, WA). The tip of the electrode is stripped for a distance of approximately 1 mm. The rear connector end of the electrode wire was stripped for a distance of about 5 mm and was threaded through a stainless steel hypodermic tube that in turn was threaded through a polyimide tube. The wire was glued into the tubing with cyanoacrylate so that the one mm uninsulated on the lead end of the wire protruded from the tubing.

### Behavior

Monkeys were trained for appetitive food pellets. A touchscreen was enclosed in a metal enclosure that also included a computer and a pellet dispenser. The training apparatus was mounted on the cage. Custom software ran 96 behavioral trials per day. In the working memory task, three rectangles were outlined on the screen, one centered and high on the screen, and two lower squares to the left and right of each other. The top rectangle illuminated with the cue color. Upon being touched, the square returned to the background color. The three rectangles were background color during the delay period. After the delay ended, the two lower rectangles were illuminated with the cue color, and a distractor color, until the monkey’s choice by touch. Successful trials triggered the release of a food pellet, and unsuccessful trials triggered a time-out. Three successful trials resulted in a longer delay period, approximately a 50% increase each time. One unsuccessful trial resulted in a decrease in trial length to the previous shorter trial length, which was roughly 33% shorter. This three-up, one-down adaptive tracking can be used to measure the 79% threshold by averaging an even number of trial reversals(13). The reversals are trials in which upward adaptive tracking changes to downward tracking, or vice versa. All animals were started on short intervals and tracked to their working range over the early trials. Only reversals after trial 30 were used for statistical analysis, and as many reversals as possible were used provided an even number were available. Only sessions in which greater than 70 of the 96 daily trials were used, and animals were used for a behavioral assessment only if they completed 70 trials in the preponderance of the sessions used for that assessment. This cutoff was used to avoid biases based on the time of session, and because it would result in too few reversals for adequate rigor in analysis. The same task was used each day. The colors used were generally yellow, blue, and red, but in some cases if animals had very long thresholds, a color scheme with more similar colors, such as three shades of blue, were used to keep the trial lengths shorter. The same colors were used, per animal, for all data presented.

### Behavioral statistics

The statistical approach for behaviors used the mean of two adjacent adaptive tracking reversals as the primary measure. A tracking reversal occurs when the adaptive tracking algorithm changes from upward tracking to downward tracking, or vice versa, provided the delay being used has not been at the lowest allowed value more than 3 trials. If the number of tracking reversals on any day was an odd number, the earliest tracking reversal that day was dropped to ensure that each downward tracking reversal had a paired upward tracking reversal. The expected value of the mean of each pair of reversals using 3-up 1-down tracking is the 79.4% threshold (13). A 96 trial behavior included from 3 to 8 adaptive tracking reversal pairs. As the tracking reversals were in a geometric, rather than an arithmetic, progression, the logarithm of the delay of each reversal was taken prior to statistical testing to approximate normality. Primary statistical tests were generalized linear mixed-effect model regressions. Each monkey was treated as a random effect. Stimulation or drug condition was treated as a fixed effect. Significant results from convergent models were followed by posthoc t-tests, provided the p values for significance met criteria established by the false discovery rate method. Software for statistics were from the python statsmodels and pingouin libraries.

### Donepezil administration

Donepezil (Sigma-Aldrich, St. Louis, MO) was administered after behavioral training each day at a dose of 100-200 μg/kg per day. The 200 μg/kg dose was used for each animal, but in some animals, motivation for the behavior was reduced presumably from appetitive interactions, and lower doses (but above 100) were used in those animals. The highest dose tolerable, up to 200, was always used. The donepezil was delivered in a date after each behavioral session. The rationale for this dose was based on previous behavioral and functional brain imaging data (dose range 0.05-0.250 mg/kg) in rhesus monkeys as well as to approximate the typical (oral) dose range used in human patients(58).

### Electrode locations in positive controls

The mediolateral center of the globus pallidus floor, 2 mm posterior to the crossing of the pallidus by the anterior commissure, was the target in implantation. This target was chosen to avoid stimulating three off-target white matter tracts, the optic nerve, the anterior commissure, and the internal capsule. In three of the animals, the electrodes missed their targets as confirmed by imaging or post-mortem examination. In two of the cases, the midline between the ear canals in the stereotaxic device was more than 5 mm lateralized relative to the brain midline. These three animals constitute the positive stimulation controls in Figure S3.

### Stimulation

All stimulation was delivered by spinal cord stimulators (Spectra WaveWriter, Boston Scientific) programmed to deliver bipolar pulses and externally controlled by the clinician programmer. Animals that had two electrodes appropriately positioned, and registering impedances in range, finished behavioral trials most days, and were not scheduled for the PET study, received bilateral stimulation after unilateral stimulation. The number meeting these criteria were not adequate to assess bilateral vs unilateral impacts. We estimate the activating function radius using the Stoney square root law(59) as 700 microns multiplied by the square root of the ratio of the current to 0.500 mA over a 100 μS phase.

### FDG-PET image acquisition and preprocessing

All PET procedures were performed at the Vanderbilt site. Three animals in the treated group with 9.5 ± 2.7 kg average body weight were each imaged once prior to bilateral electrode implantation and once during or shortly after receiving unilateral NB stimulation (2^nd^ scan 288 ± 69 days (M ± SD) after 1^st^ scan, 85 ± 30 days after stimulation onset). The animal was anesthetized with ketamine (15 -20 mg/kg) for induction and isoflurane (1 - 5%) for maintenance. The animal received an intravenous injection of [^18^F]Fluorodeoxyglucose (FDG) with an average total radiopharmaceutical dose of 173.5 ± 5.1 MBq, before 31 frames of positron emission tomography (PET) scans were acquired between 2 min and 161 min on average after tracer injection. PET scans with a voxel size of 1×1×1mm were acquired with a Philips Vereos PET/CT scanner.

PET scans were preprocessed using the “volreg” and “align” blocks of the afni_proc.py pipeline in AFNI. Each sequence was motion-corrected, aligned to the T1w image using a rigid-body transform and further mapped to the standard template space using the transforms generated by @animal_warper. The final mapping accuracy was verified visually. Standard uptake values (SUV) were then computed for each frame using equation 3, where*r_t_* is the radioactivity activity concentration (Bq/mL) of the image acquired at time *t* (s) since tracer injection, *w* is the animal’s body weight (g), *d* is the total injected radiopharmaceutical dose (Bq) and *T*_1/2_ is the half-life (s) of FDG.

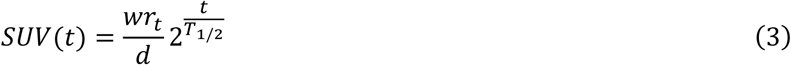

SUV values of frames taken between the 40^th^ and 60^th^ min(60) in each sequence were averaged across frames and then normalized to the mean SUV in the ipsilateral pons due to its low age-related changes(61) and its lack of cholinergic projections from the basal forebrain. Normalizing within each hemisphere controls for any potential lateralized differences in signal intensity not specific to brain tissue. This procedure generated the Standard uptake value ratios (SUVr) measure which served as the dependent variable in subsequent analyses. The normalization with the ipsilateral pons is compared to a more typical cerebellar normalization in Figure S5.

### PET SUVr data analysis

To study the impact of stimulation on SUVr in the ipsilateral hemisphere, we utilized a linear mixed-effect model using MATLAB function “fitglme” by including “treatment stage”, “stimulated side” and their interactions as fixed-effect terms along with “subject” as a random-effect intercept term (equation 4). The reference levels of “treatment stage” and “stimulated side” were the baseline period and the contralateral hemisphere respectively. Thus, a positive effect of stimulation on SUVr in the ipsilateral hemisphere would be indicated by a positive coefficient for the interaction term, while a negative coefficient for the main effect of “treatment stage” would indicate an age-dependent decrease in SUVr orthogonal to stimulation.

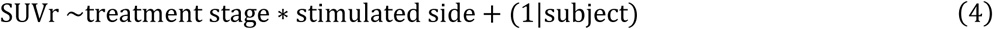

For voxel-wise analysis, this model was computed for each voxel, resulting in 12 observations for each voxel and thus degrees of freedom (8, 1) for each fixed effect. For an effect to be significant in a voxel, the voxel-level one-tailed alpha level was chosen to be 0.005 (equivalent to a 0.01 two-tailed test). To control for familywise Type I error rate(62), we first estimated the spatial noise autocorrelation function (ACF) from each unilaterally-masked, individual SUV timeseries using AFNI’s 3dFWHMx function which assumed a mixture of Gaussian and exponential functions. The anatomical correlation was removed by fitting the timeseries within each voxel with a set of 5 reference functions (constant, linear, quadratic, sine, and cosine) using the “-detrend” option. Geometric means of the ACF parameters from all individual timeseries make up the final population average ACF (equation 5) with respect the Euclidean distance between voxels *r* in units of mm.

Then using AFNI’s 3dClustSim function, we simulated the null distribution where each voxel in the unilateral brain mask was only significant by chance due to noise with spatial correlation described by the population average ACF. A spatially connected (first nearest neighbor, using the “-NN 1” option) cluster of all positive or all negative significant voxels (using the “bi-sided” output) larger than 72.59 mm^3^ (4646 voxels) in size was found to be less likely than 0.05 in the null distribution. For each fixed effect in the model, clusters larger than this threshold was considered significant. The regions encompassed by the significant clusters were manually identified by overlaying the clusters on the template along with the CHARM and SARM atlases in 3D Slicer. For surface visualization, the T values within significant clusters were projected to the white matter surface of the NMT v2.0 template by averaging between the white matter surface and the pia surface using AFNI’s 3dVol2Surf function. The output was rendered using SUMA(63).

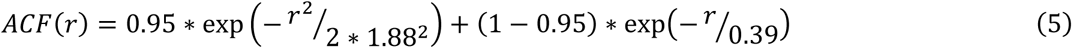

For ROIs-based analysis, we followed the brain parcellation by the 2^nd^ level of the CHARM atlas which segments the neocortex into 17 regions. SUVr was averaged within each unilateral ROI ipsilateral and contralateral to each subject’s stimulation side respectively. An additional random intercept for ROIs were added to the GLMM to test for average changes across the whole neocortex.

### Tissue plasminogen activator experiments

The tissue plasminogen activator (tPA) assay used was AbCam ab108914. Cerebrospinal fluid (CSF) samples were selected from lumbar puncture except for two animals implanted with Ommayas, in which case the CSF was sampled from the Ommaya (Medtronic CSF ventricular reservoir 24106a, neonatal sizing). All CSF samples for tPA analysis were collected and analyzed at the Augusta location.

## Supplementary Figures

**Figure S1.**
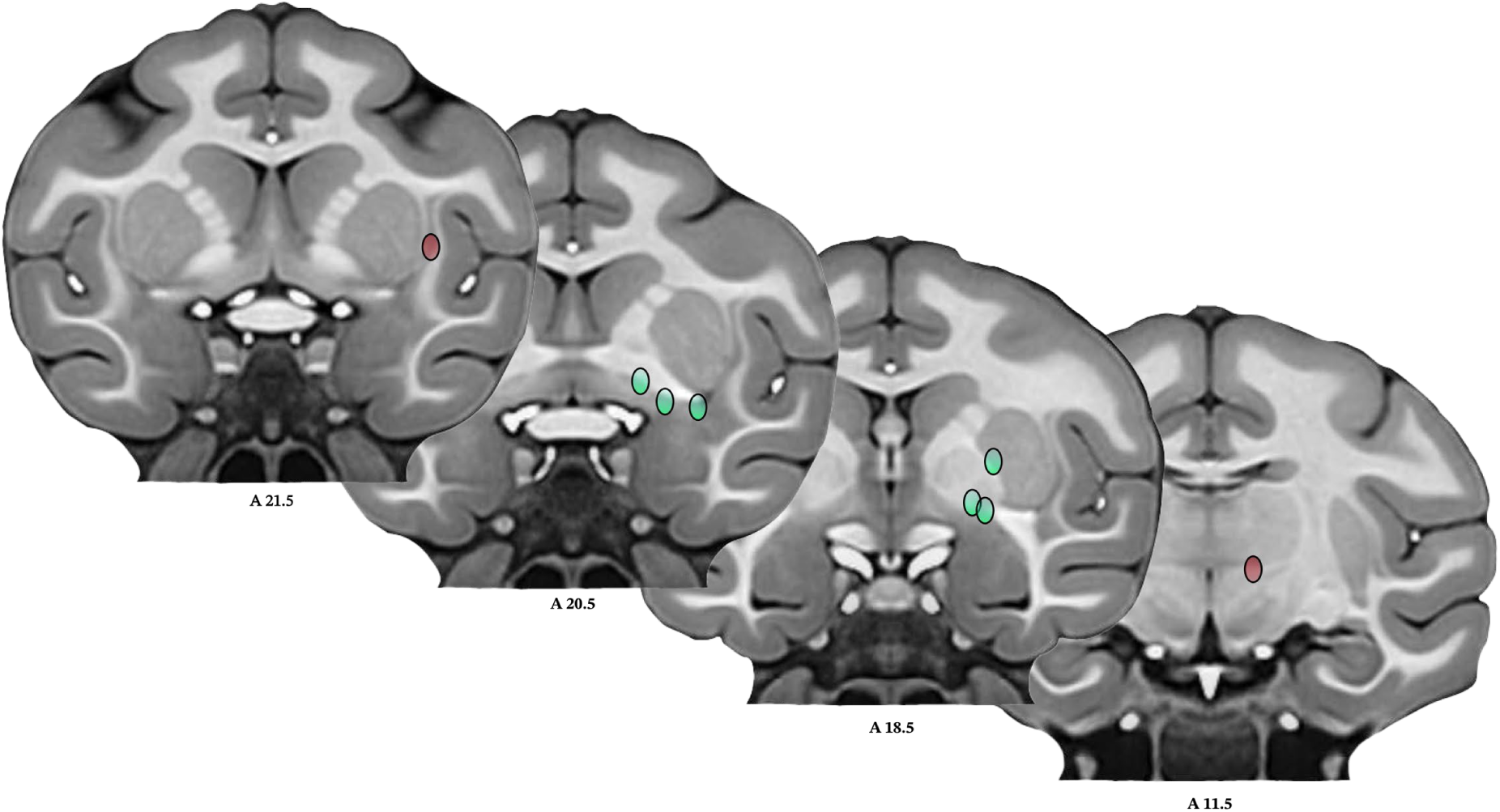
Electrode tip positions in monkeys recovered by overlaying CT and MRI, or by electrolytic lesion localization. Blue indicates recovered positions considered in the inferior globus pallidus or nucleus basalis of Meynert/substantia innominata. Red indicates electrode tips that were not considered placed correctly for study (positive stimulation controls in Figure S2). All positions are mirrored into the same hemisphere for visualization. Sections are taken from the NMT template V2 with A-P coordinates indicated. The shape of the tip positions was determined by using a 0.75 mm radius around a 1 mm line, to approximate the activating function surrounding our 1 mm wire exposure on the electrode.

**Figure S2.**
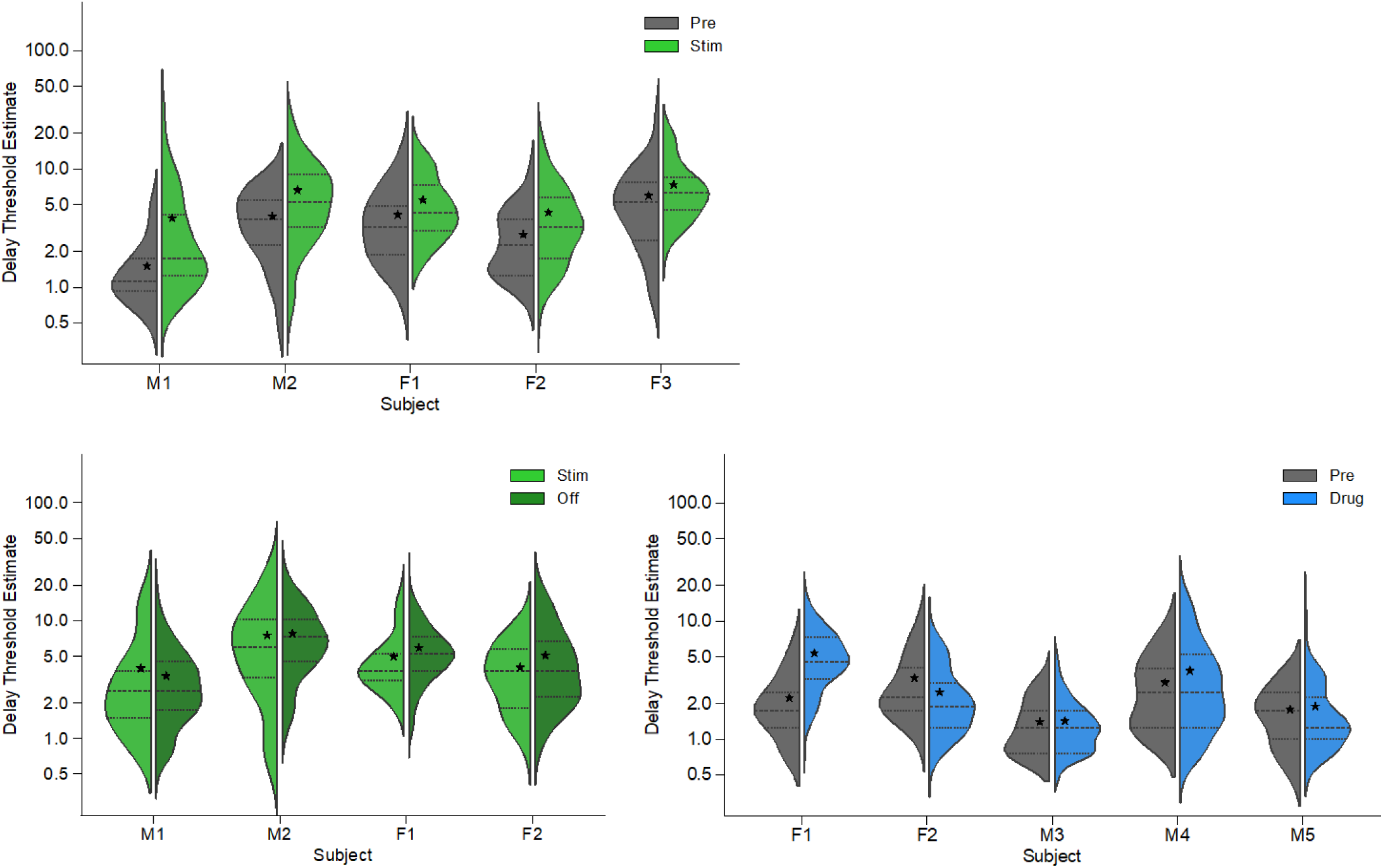
Split violin plots detailing all data points for averages in Fig. 1. Top Left indicates data collected during Pre vs. Stimulation conditions, also shown in Fig. 1D. Bottom Left indicates data collected during Stimulation vs. Off conditions, also shown in Fig. 1F. Bottom Right indicates data collected during Pre vs. Drug conditions, also shown in Fig. 1G.

**Figure S3.**
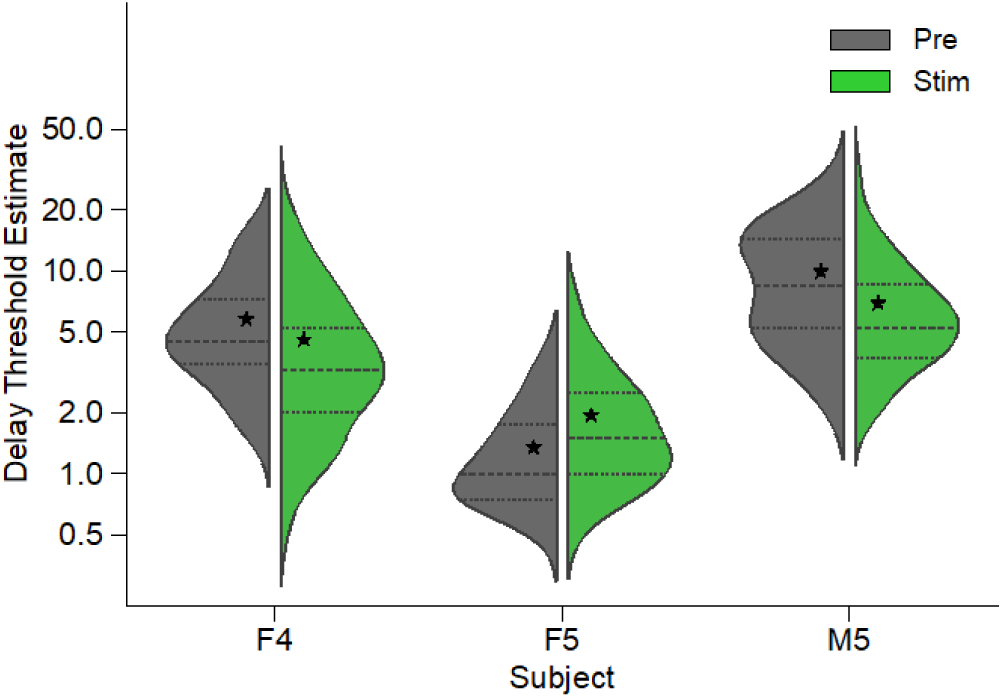
Split violin plots of adaptive tracking reversal delay pair means for three animals that were implanted off-target, and received stimulation from electrodes not placed in or below the inferior globus pallidus.

**Figure S4.**
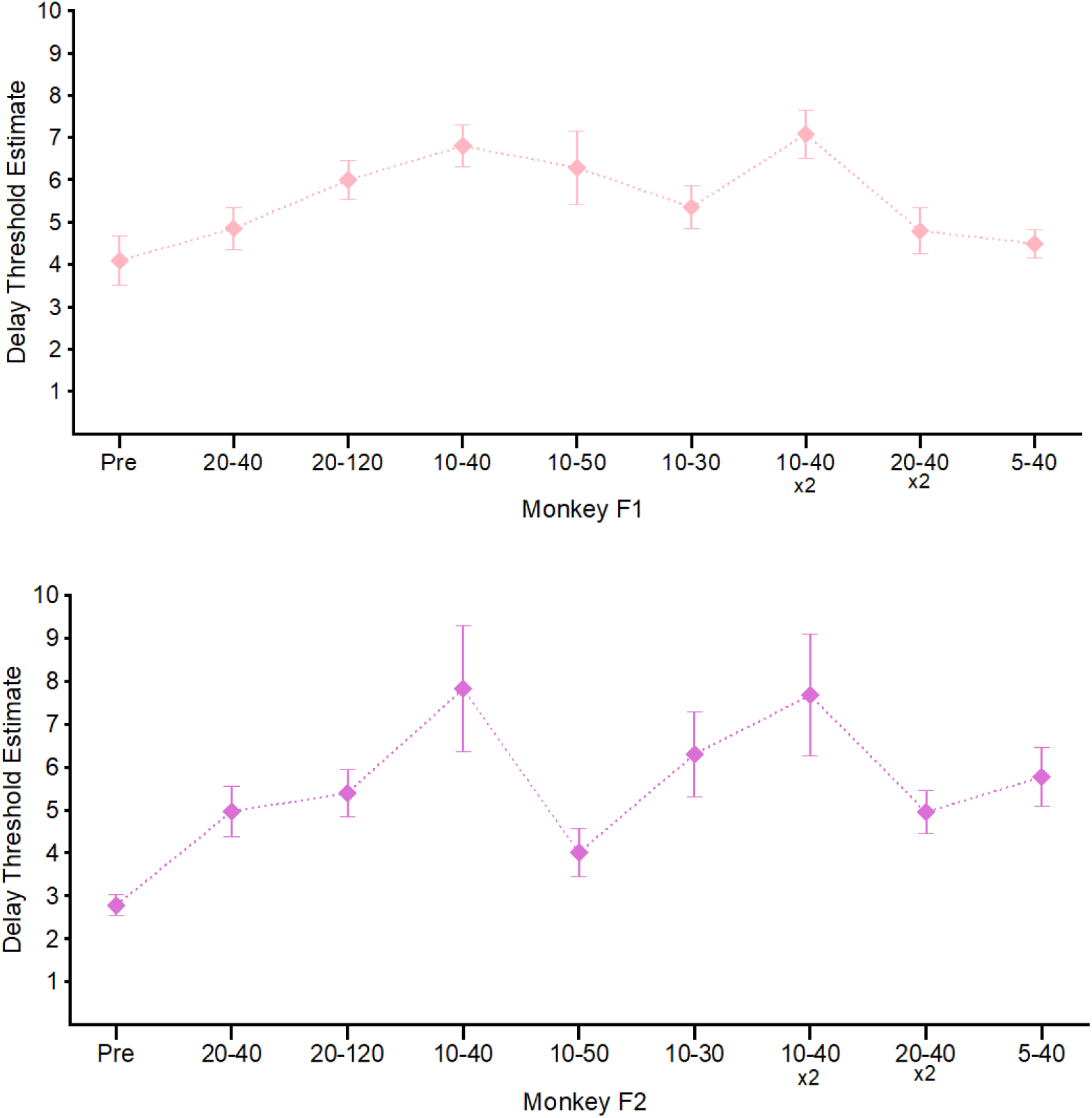
Delay threshold estimates for two animals as a function of stimulation pattern. Data points from left to right were collected in chronological order. Four to five behavioral sessions were performed at each pattern. Means and standard errors are marked in each plot.

**Figure S5.**
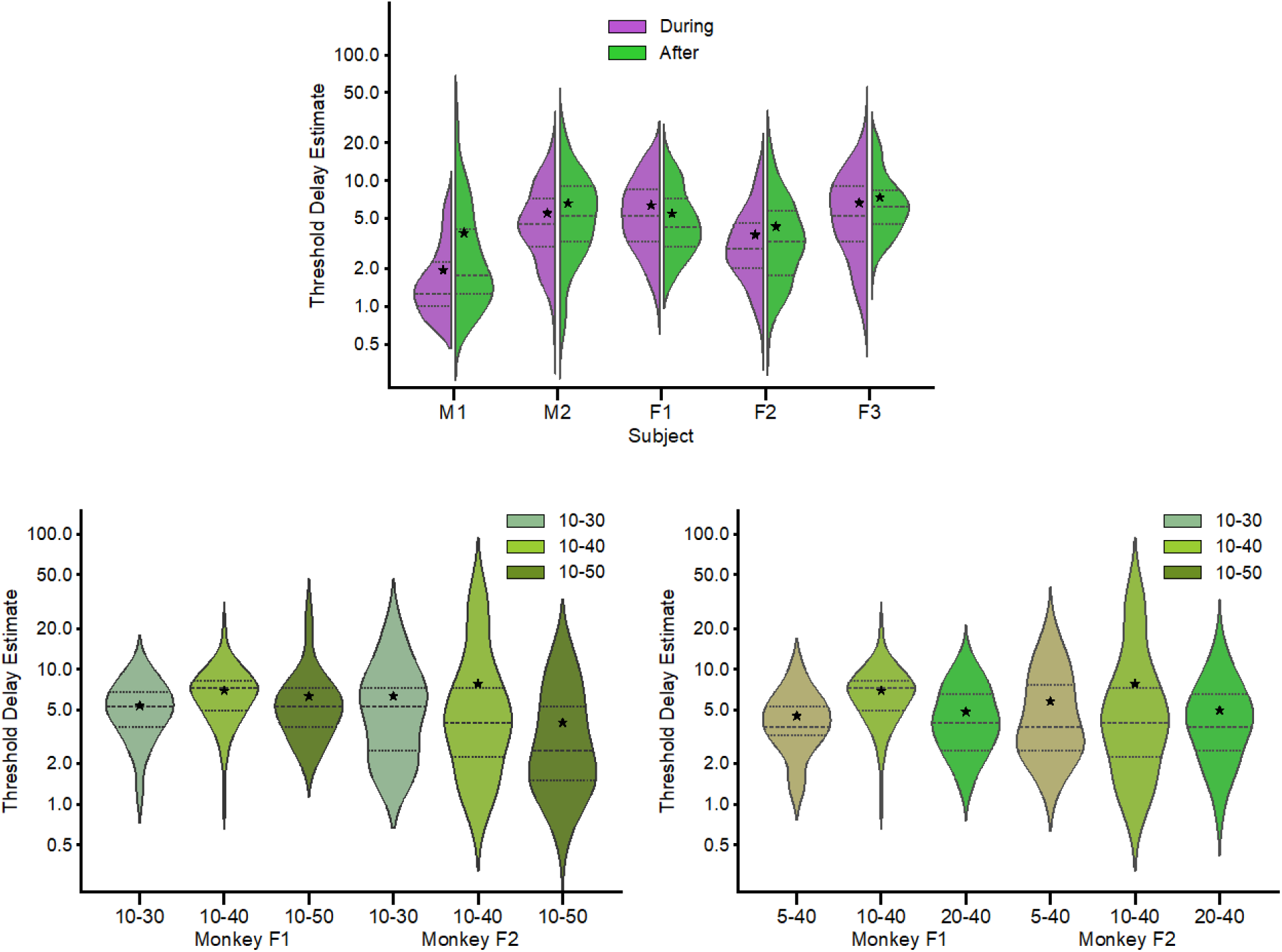
Violin plots detailing all data points for averages seen in Fig. 3. Top indicates data collected during a period when monkeys received stimulation each day, either during or after behavior task performance. Bottom Left indicates data collected during tuning of the Recovery parameter of stimulation. Bottom Right indicates data collected during tuning of the Stimulation parameter.

**Figure S6.**
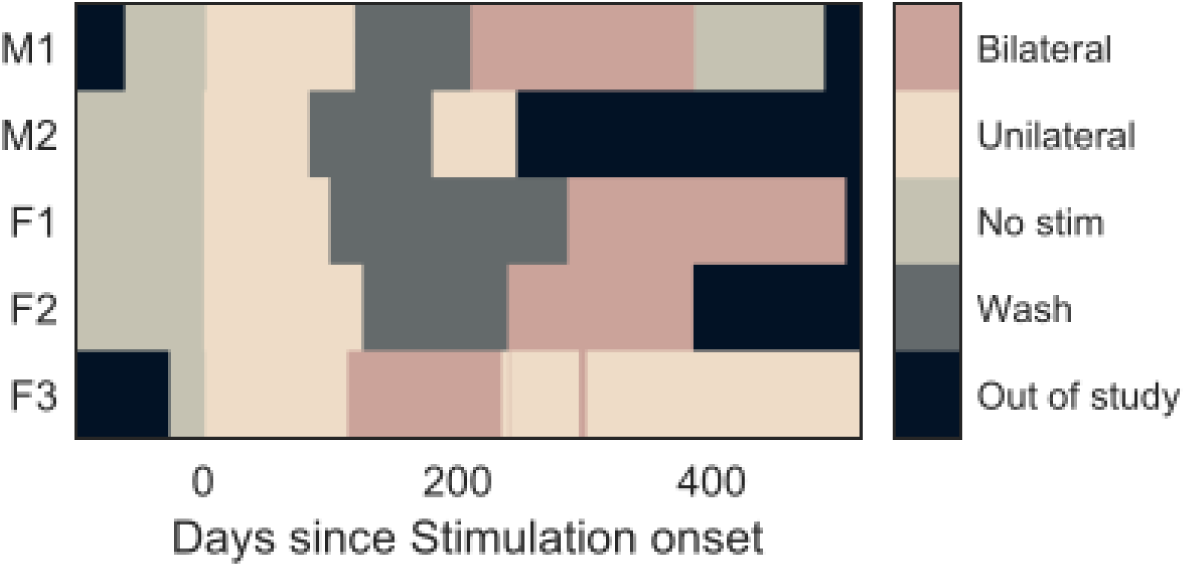
Stimulation history for animals in Figure 1D-F, and Figure 2. All animals received an 8-12 week period of unilateral stimulation beginning at day zero. Four had periods of at least 12 weeks without stimulation after the initial stimulation period. Three of the animals had bilateral working electrodes and received bilateral stimulation. Two of those are shown in Figure 3B-C.

**Figure S7.**
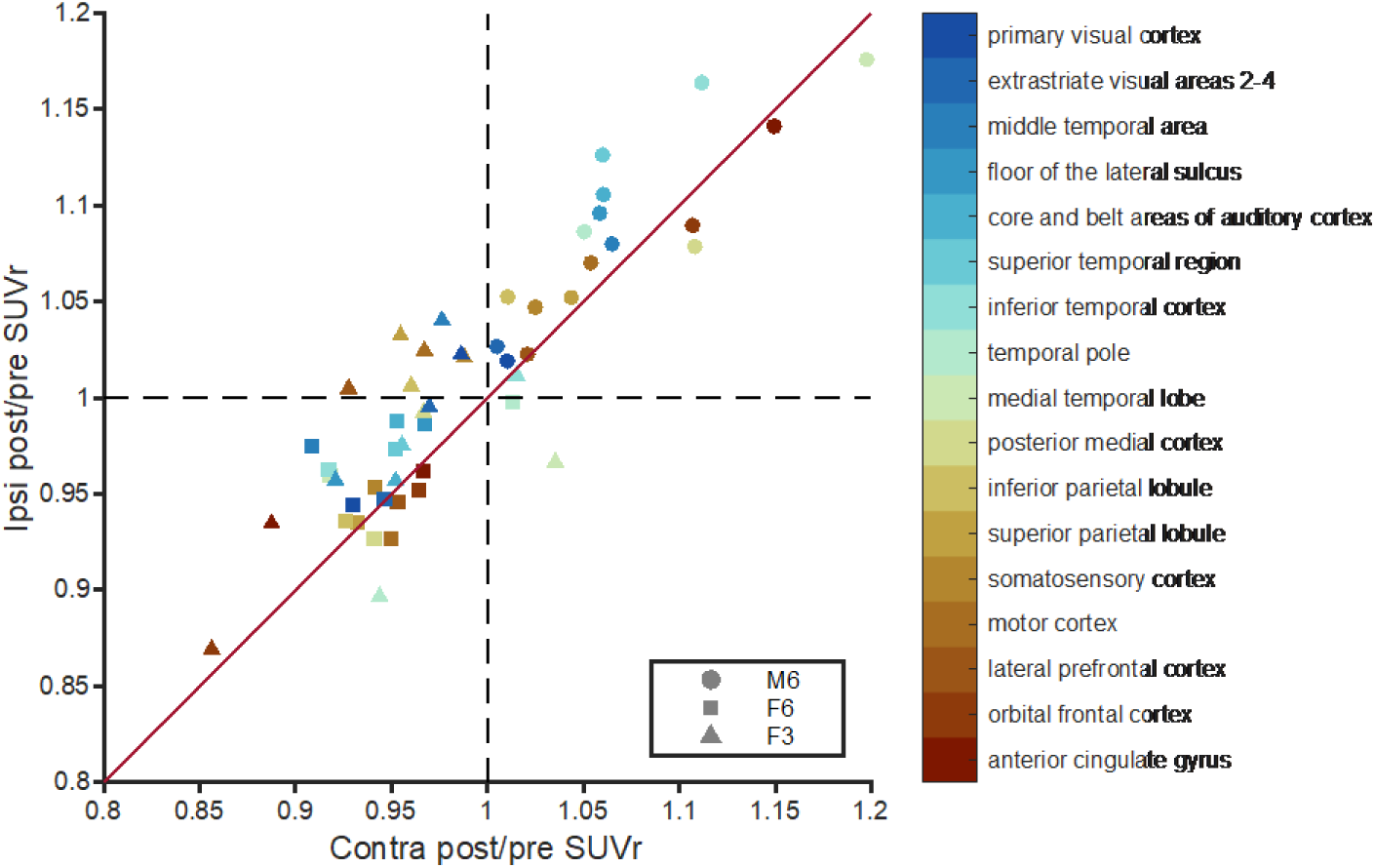
Comparison of 12 week PET SUVr change ratios between stimulated and unstimulated hemispheres normalized by the cerebellum signal in 17 cortical regions in three animals. Findings complement those with pons normalization in Figure 4. The Ipsi hemisphere received stimulation, and the Contra hemisphere did not.

